# IBDkb: an AI-enhanced integrative knowledge base for inflammatory bowel disease research and drug discovery

**DOI:** 10.64898/2026.01.27.702180

**Authors:** Liwen Tao, Shiqing Shi, Ruixin Zhu, Zhuo Liu, Bin Yang, Lei Liu, Wanning Chen, Qian Long, Na Jiao, Guoqing Zhang, Pingping Xu, Dingfeng Wu

## Abstract

IBDkb (Inflammatory Bowel Disease Knowledge Base; https://www.biosino.org/ibdkb) is a freely accessible, integrated web-based platform that systematically curates and harmonizes multi-source data related to inflammatory bowel disease (IBD). To address the fragmentation and therapeutic gaps in existing specialized resources, IBDkb establishes a unified framework featuring advanced full-text search, interactive visualizations, cross-module knowledge graphs, and AI-powered utilities for real-time literature retrieval, trend analysis, text/PDF interpretation, and domain-specific conversational assistance. The platform currently integrates 98,453 research articles, 3,390 clinical trials, 200 investigational drugs, 200,606 bioactive compounds, 103 therapeutic targets, 77 experimental models, 12 pathogenesis summaries, and 15 treatment strategies. These integrated tools facilitate efficient exploration of complex associations among drugs, targets, trials, and mechanisms, thereby accelerating hypothesis generation and translational research in IBD. The platform is openly available without registration and supports data downloads. A case study on structure-aware drug comparison further demonstrates its utility in facilitating cross-disease drug repositioning hypotheses.

## Introduction

Inflammatory bowel disease (IBD) is a chronic and complex inflammatory disorder of the gastrointestinal tract, comprising two main conditions: Crohn’s disease (CD) and ulcerative colitis (UC) [1]. It is characterized by a lifelong, relapsing-remitting course with symptoms such as diarrhea and abdominal pain [2, 3], and is associated with an increased risk of colorectal cancer [4]. The necessity for long-term treatment and ongoing clinical monitoring substantially impairs patients’ quality of life. Owing to its increasing incidence and prevalence worldwide, IBD now affects millions and represents a growing global public health challenge with substantial clinical and socioeconomic burdens [5, 6].

IBD arises from a multifactorial etiology involving intricate interactions among genetic susceptibility, environmental exposures, immune dysregulation, and alterations in the gut microbiota [7]. Traditional management has relied on broad anti-inflammatory and immunosuppressive therapies, including aminosalicylates, corticosteroids, occasionally thiopurines and antibiotics for mild to moderate IBD. For moderate to severe IBD, the introduction of biologic agents such as anti-tumor necrosis factor (TNF) monoclonal antibodies has revolutionized IBD treatment [1]. However, treatment efficacy remains limited; for instance, only approximately 30-50% of patients achieve clinical and mucosal remission with anti-TNF therapy. Consequently, emerging therapeutic strategies are being explored to more precisely target IBD pathogenesis. These include novel biologics (e.g. anti-IL-12/23, anti-α4β7/αEβ7 agents), small-molecule inhibitors such as Janus kinase (JAK) inhibitors, as well as cell- and microbiome-based therapies [8]. Collectively, these advances underscore the shift toward precision medicine in IBD.

Concurrently, advances in high-throughput technologies and clinical informatics have generated increasingly complex and multi-dimensional datasets in IBD research, evolving from single-omics profiles to integrative multi-omics frameworks. Study designs have also expanded from single-center cohorts to multi-center cohorts and specialized populations [9–11]. While these advances have greatly expanded the scope of IBD research, they have also resulted in substantial data heterogeneity and fragmentation across studies and data types. To address these challenges, several specialized databases have been developed, such as the IBD Exomes Browser for genetic variant exploration [12], the Inflammatory Bowel Disease Multi’omics Database (IBDMDB) [13], the Inflammatory Bowel Diseases Integrated Resources Portal (IBDIRP) [14], both of which focus on multi-omics data, the Ulcerative Colitis Database (UCDB) [15], the IBD database (IBDDB) [16], and the IBDB [17], which curate IBD-associated genes, as well as transcriptome-centered resources such as IBDTransDB [18] and scIBD for single-cell meta-analyses [19].

Despite their individual utility, these resources differ markedly in scope, update frequency, data granularity, and analytical functionality. Crucially, most are primarily descriptive and omics-centric, with limited support for integrating therapeutic knowledge, such as systematic curation of IBD-related drugs, treatment regimens, clinical trial evidence, experimental models, and drug-target relationships. Standardized integration and cross-linking among genomic findings, clinical trials, therapeutic strategies, and drug development-relevant resources remain largely lacking.

This fragmentation hinders data interoperability, sharing and downstream reuse, particularly in the context of IBD drug discovery, mechanism-driven therapeutic development, and AI-assisted drug design [20]. Therefore, there is an urgent need for an integrated, multi-level knowledge base that bridges molecular mechanisms, pharmacological interventions, clinical evidence, and experimental models to directly support IBD therapeutic research and drug development pipelines.

To bridge these gaps, we developed IBDkb, a comprehensive web-based knowledge base that integrates multi-source IBD-related data with manual curation, advanced visualization, and AI-enhanced tools to support translational research and drug development.

## Materials and Methods

### Dataset collection and curation

IBDkb integrates data from diverse public sources, including research articles, clinical trials, drug information, therapeutic targets, bioactive compounds, and experimental models (Figure 1A). As of January 10, 2026, IBDkb comprises 98,453 annotated research articles, 3,390 clinical trials, 200 investigational drugs, 1,404 drug targets, 200,606 bioactive compounds, 77 experimental models, 103 IBD-specific therapeutic targets, 12 pathogenesis records, and 15 therapeutic strategy records. Literature records were retrieved from PubMed using keyword-based searches and automated queries via the Python Entrez toolkit. The search strategy focused on articles published within the past two decades, employing core terms including “inflammatory bowel disease”, “Crohn’s disease”, and “ulcerative colitis” as well as their relevant synonyms and MeSH terms. For each publication, the title, abstract, PubMed ID (PMID), journal name, and publication date were extracted. Clinical trial information was collected from internationally recognized registries, including ClinicalTrials.gov (https://clinicaltrials.gov), the World Health Organization’s International Clinical Trials Registry Platform (https://trialsearch.who.int), and DrugBank [21], covering essential details such as study design, recruitment status, enrollment criteria, and trial phase. From the collected research articles, IBD-related therapeutic strategies, pathogenesis, and associated diseases were manually extracted. Investigational drugs were compiled from published studies, registered clinical trials, and the DrugBank database. Drug-target relationships were derived from DrugBank [21] and PubChem [22], and bioactive compounds were retrieved from ChEMBL [23]. Data on therapeutic targets were gathered from the Online Mendelian Inheritance in Man (OMIM) database [24], the Therapeutic Target Database (TTD) [25], and UniProt [26]. Information on experimental models, pathogenesis, and therapeutic strategies were manually extracted and annotated from key publications.

**Figure 1.**
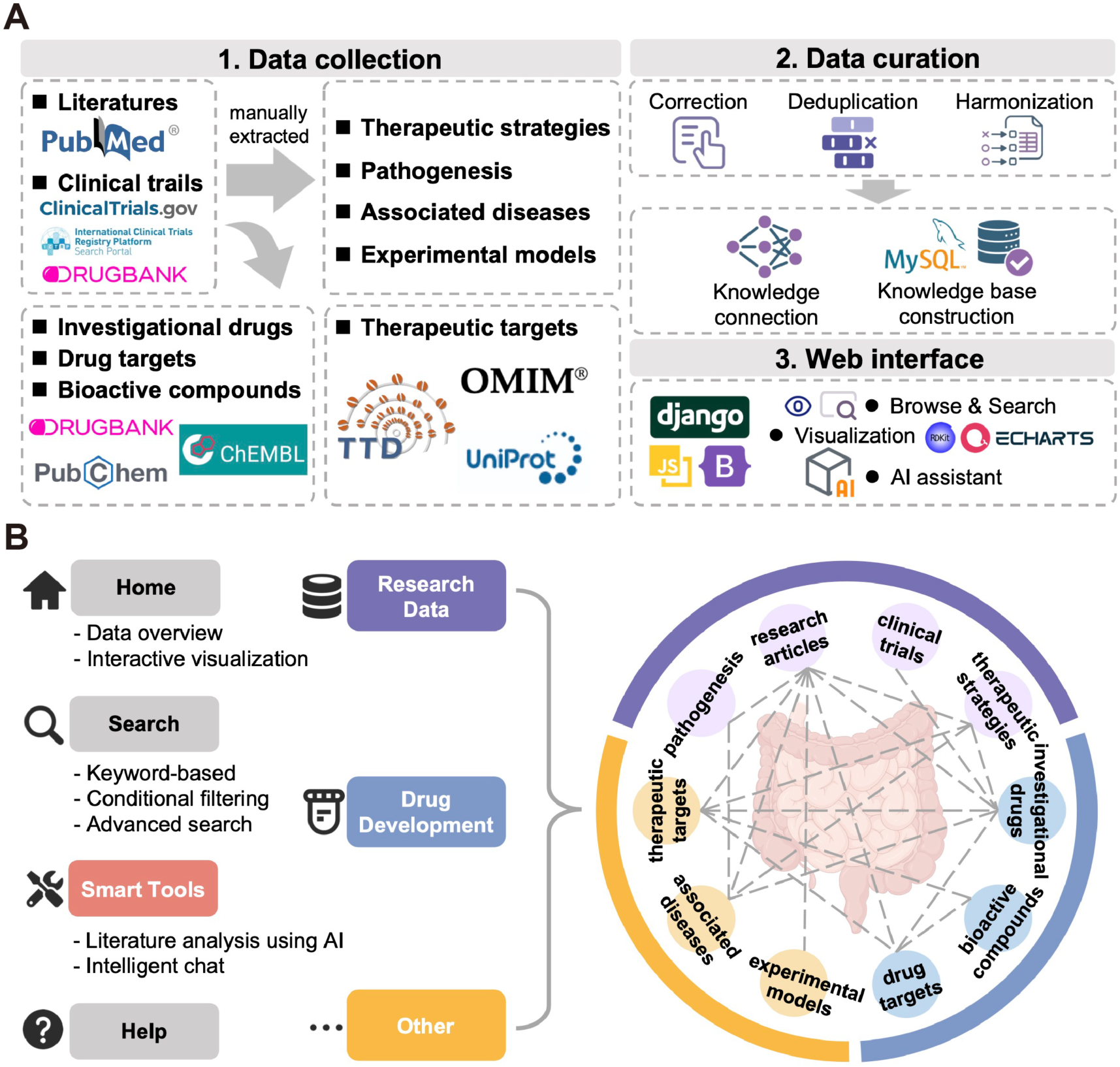
The framework and available function modules of IBDkb. (A) Schematic workflow illustrating the construction of the IBDkb knowledge base. (B) System architecture and core functionalities of the IBDkb platform.

To ensure data quality and consistency, we implemented a multi-step curation pipeline including manual correction, deduplication and format harmonization. Duplicate entries were removed using custom Python scripts based on unique identifiers and title/name matching. Ambiguous records were reviewed by domain experts. Data from heterogeneous sources were standardized into a unified structure. All datasets are periodically updated.

### Platform development and implementation

The web interface was developed using the Django framework (v5.1.2) with Python (v3.11.13), enabling a clear separation of system modules for easier feature expansion and maintenance. Data is managed using a MySQL database (v9.5.0) to support efficient storage and retrieval. The front-end uses standard Hypertext Markup Language (HTML) for content rendering, complemented by Bootstrap for responsive styling, and JavaScript for interactivity. Client-server communication is handled through asynchronous AJAX calls to the back-end API.We applied RDKit (v2023.09.1) to support chemical structure visualization for drugs and compounds. For data analytics and interactive charts, the platform incorporates ECharts (v5.4.3).

The IBDkb platform is enhanced with artificial intelligence (AI) capabilities. Specifically, large language model (LLM) ERNIE Bot (ernie-3.5-8K; Baidu, Inc) was selected as the AI engine for its strong performance in Chinese and English biomedical text comprehension and its accessible API. It powers intelligent literature analysis and conversational interaction. In addition, we developed a global IBD AI assistant based on a retrieval-augmented generation (RAG) framework, which retrieves evidence from the IBDkb knowledge corpus to provide grounded, context-aware responses. The platform is hosted on stable national servers with regular backups to ensure long-term availability. The platform further enables downloading of complete data resources, customized filtered results, and processed AI-ready datasets.

### Platform architecture and functional modules

The IBDkb platform is organized into six major sections accessible via a unified navigation bar: Homepage, Search, Research Data, Drug Development, Smart Tools, Other, and Help (Figure 1B). The homepage provides an entry point featuring a global text search bar, links to functional modules, and an interactive network visualization, a data overview panel showing temporal trends and the latest research updates.

The Research Data section contains four modules: Research Articles, Clinical Trials, Therapeutic Strategies, and Pathogenesis. The Drug Development section compiles data on Investigational Drugs, Bioactive Compounds, and Drug Targets. The “Other” section provides supplementary, curated resources, including Experimental Models, Associated Diseases, and Therapeutic Targets. Each module supports structured data display, keyword search, and customizable filtering. The Smart Tools section provides AI-assisted functionalities, including a knowledge acquisition module that retrieves published literature from PubMed Central and performs intelligent content analysis using large language models.

To enhance usability, the search page summarizes the number of matched records within each key module. Several modules support multi-attribute search, including keyword-based queries, conditional filtering, and advanced search. The help page offers a user guide, update logs, and a feedback interface.

## Results

### Overview of curated data in IBDkb

In total, IBDkb contains 98,453 research articles, 3,390 clinical trials, 12 pathogenesis records, 15 therapeutic strategy records, 200 investigational drugs, 1,404 drug targets, 200,606 bioactive compounds, 77 experimental models, and 103 IBD-specific therapeutic targets. All data are freely accessible and periodically updated via the IBDkb website (https://www.biosino.org/ibdkb/). Each module provides a concise summary view with key information (Figure 2A). Detailed information for each record is accessible via a “Details” link.

**Figure 2.**
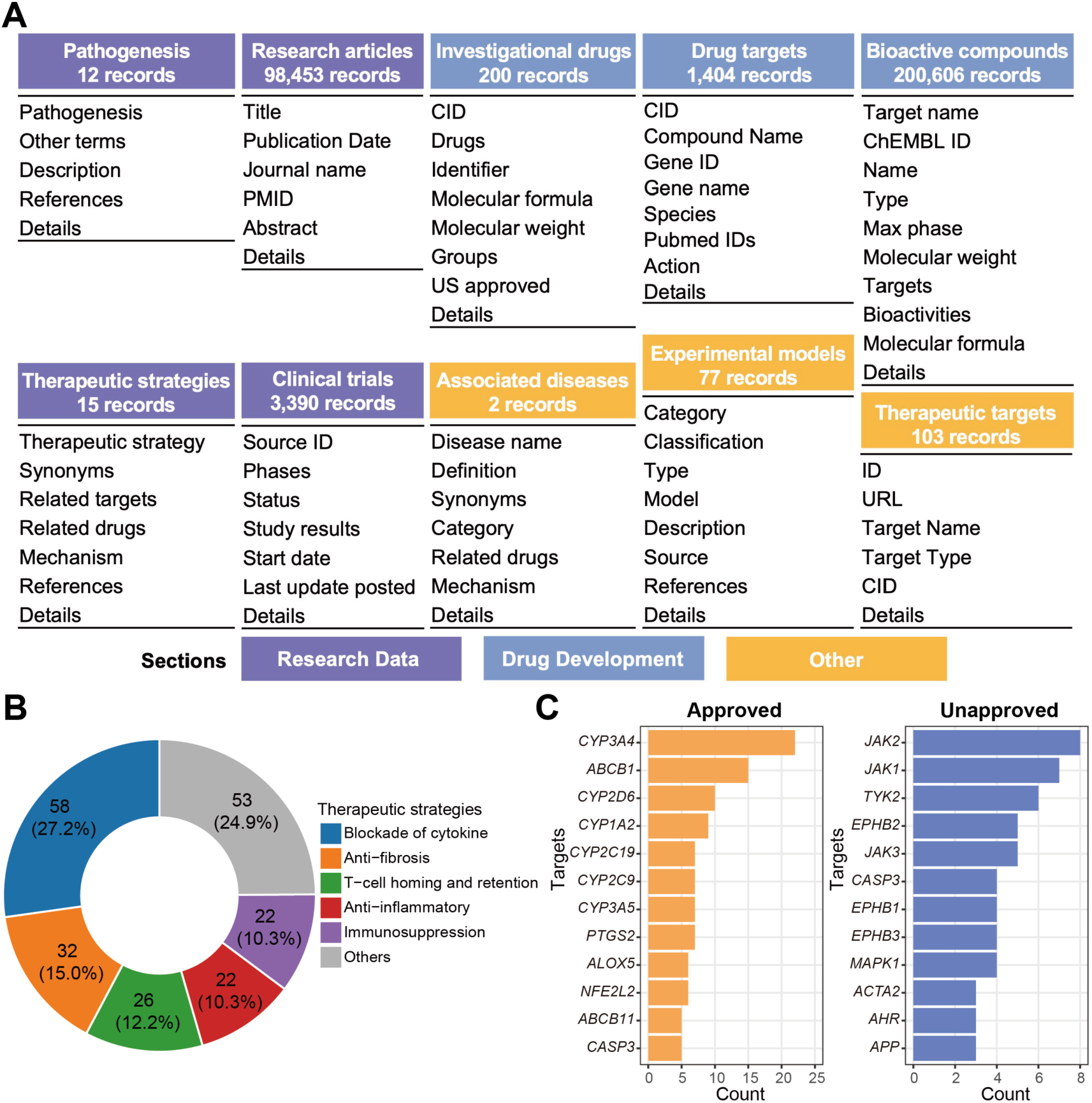
Overview of curated data in IBDkb. (A) Core information displayed in each module of IBDkb. (B) Distribution of therapeutic strategies among investigational IBD drugs. (C) Top 12 drug targets identified in approved and unapproved IBD drugs, respectively.

We summarized the therapeutic strategies of investigational drugs and observed that the majority act through cytokine signaling blockade (27.2%), followed by anti-fibrotic mechanisms (15.0%), modulation of T-cell homing and retention (12.2%), anti-inflammatory effects (10.3%), and immunosuppression (10.3%) (Figure 2B). Relatively few drugs currently target angiogenesis or stem cells transplantation, indicating that these are underexplored areas. Furthermore, we analyzed drug targets of U.S. Food and Drug Administration (FDA)-approved versus unapproved IBD. Approved drugs primarily target pathways related to xenobiotic metabolism and transport, inflammatory lipid mediator biosynthesis, and oxidative stress responses, whereas unapproved drugs are mainly enriched for targets involved in JAK-STAT signaling and intestinal epithelial homeostasis (Figure 2C).

### Platform features and utilities

IBDkb offers robust search and browsing capabilities to facilitate efficient data exploration. A convenient full-text search function allows queries across all textual data, with results organized by module and directly available for download (Figure 3A). Users can also browse data by module and apply built-in filters to refine search results. Selected modules further incorporate interactive visualizations and advanced search options to support more precise data retrieval (Figure 3B-C).

**Figure 3.**
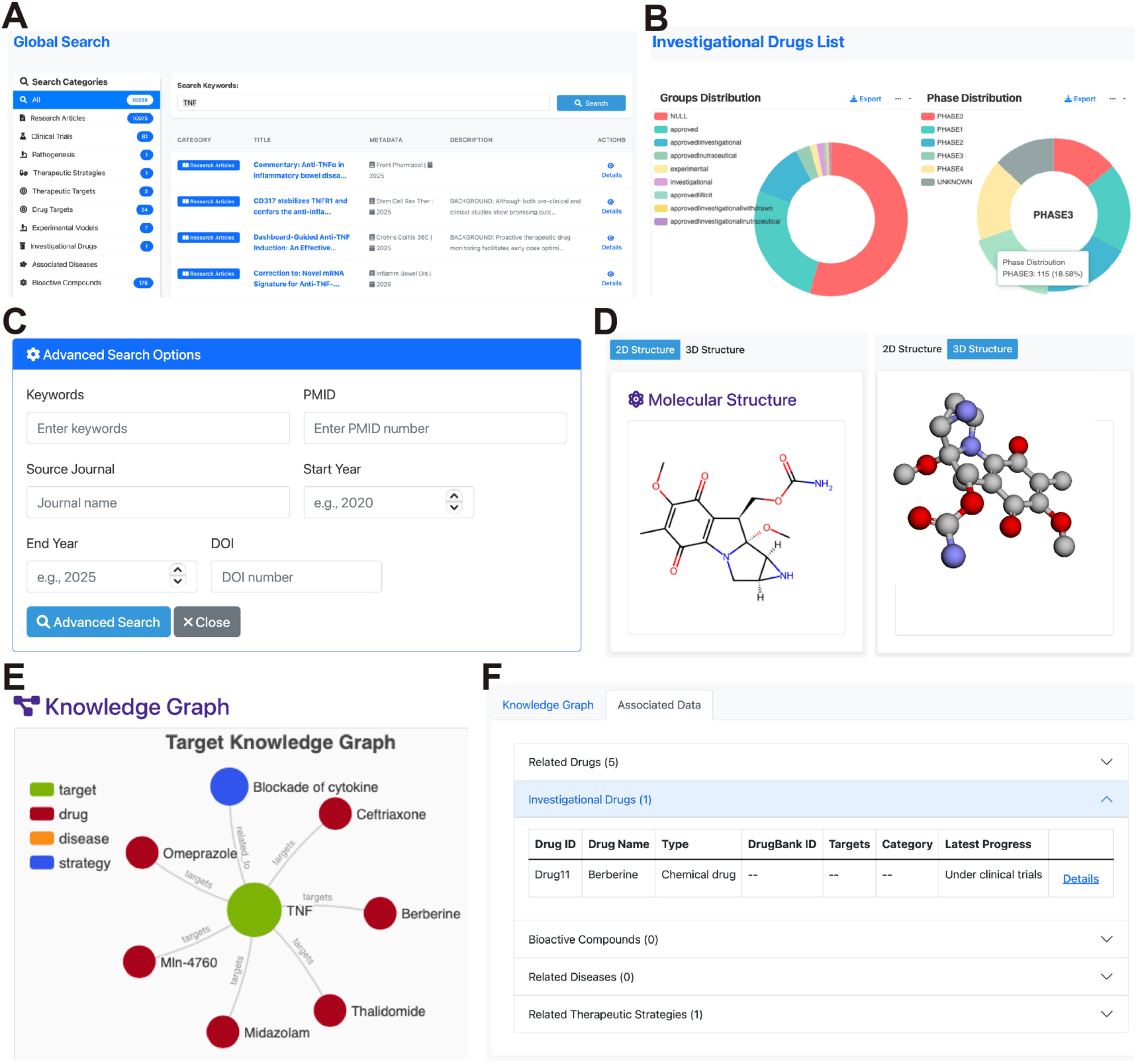
Utility of the IBDkb platform. (A) Categorized display of global search results. (B) Interactive visualization of data distribution within a selected module, exemplified by the Investigational Drugs module. (C) Advanced search functionality within the Research Articles module. (D) 2D and 3D molecular structure visualization, shown using Mitomycin A from the Bioactive Compounds module as an example. (E) Interactive network graph illustrating relationships across associated modules, exemplified by the TNF gene from the Drug Targets module. (F) Corresponding textual information displayed alongside the network, exemplified by the TNF gene from the Drug Targets module.

To ensure transparency, IBDkb provides direct links to original sources. Pathogenesis and therapeutic strategy record is accompanied by a list of related literature references, so users can obtain more detailed background information from those cited studies. The Drug Development section integrates external resources and visualization tools. Drug-related record includes links to relevant clinical trials or literature. The Investigational Drugs and Bioactive Compounds modules provide 2D and 3D molecular structure visualizations (Figure 3D). Notably, IBDkb highlights cross-module relationships (Figure S1A). In the detailed view of Investigational Drugs and Drug Targets modules, users can see connections to other modules (e.g., associated clinical trials, literature, or therapeutic strategies) via interactive network graph and text descriptions (Figure 3E-F).

In the “Other” section, the Experimental Models and Associated Disease modules provide reference links for each entry, enabling users to directly access supporting information. Similarly, the Therapeutic Targets module offers cross-module association details. In addition, IBDkb provides downloadable resources, including curated datasets and AI-ready data comprising MolFormer-based embeddings and molecular fingerprints for drugs and bioactive compounds (Figure S1B). These AI-ready datasets can be directly applied to artificial intelligence and machine learning models, enabling efficient and scalable analyses for disease-related drug discovery and knowledge exploration.

### AI-assisted knowledge acquisition module

The Knowledge Acquisition module in the Smart Tools section enhance research capabilities through three main functionalities: literature statistics, literature analysis, and an intelligent conversational interface.

This module integrates with the PubMed Central for real-time retrieval of the latest literature by keyword, filterable by year range. It provides trend analysis visualizations of publications counts over time and average annual counts (Figure 4A). A journal distribution visualization shows how results are spread across academic journals (Figure 4B).

**Figure 4.**
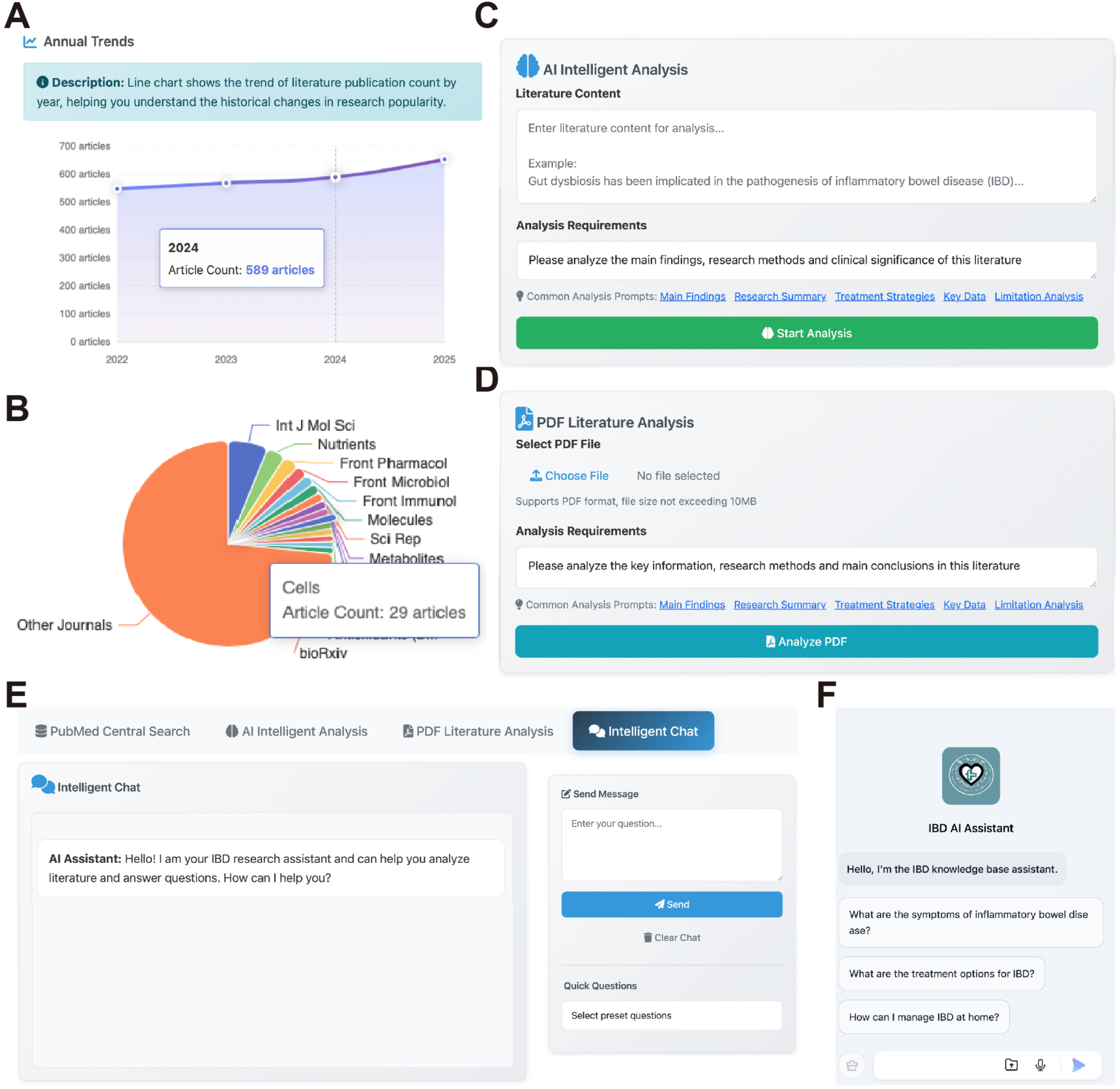
AI-assisted knowledge acquisition in IBDkb. (A) Line chart showing the annual publication trend (2022-2025) for the search term “5-Hydroxytryptophan”. (B) Pie chart illustrating the journal distribution of literature in the same search results (2022-2025, “5-Hydroxytryptophan”). (C) AI-based intelligent analysis of user-input literature content. (D) AI-based intelligent analysis of uploaded PDF literature files. (E) Intelligent chat function for interactive querying and contextual reasoning. (F) Global AI assistant accessible across the platform.

The AI-driven literature analysis component supports both direct text input and PDF upload, enabling intelligent analysis based on user-defined requirements (Figure 4C-D). To further facilitate knowledge acquisition, the module incorporates an intelligent conversational interface that supports continuous, context-aware dialogue (Figure 4E). Additionally, a global RAG-based AI assistant is accessible throughout the platform, providing expert-level guidance and enhancing the overall research experience (Figure 4F).

### Case Study: Structure-Aware Drug Comparison and Repositioning

To demonstrate IBDkb’s utility for integrative drug-centric knowledge discovery, we conducted a comparative case study focusing on investigational drugs for IBD and non-alcoholic fatty liver disease (NAFLD) (Figure 5A). IBD drugs were retrieved from IBDkb, while NAFLD-related drugs were obtained from NAFLDkb [27]. The curated datasets enabled direct input into large-scale language models. Molecular representations were generated using the MolFormer large-scale chemical language model [28], followed by UMAP dimensionality reduction with n_neighbors=15 and min_dist=0.1 for visualization (Figure 5B, Table S1). Drugs associated with IBD and NAFLD formed two well-separated clusters, indicating disease-specific chemical characteristics were captured.

**Figure 5.**
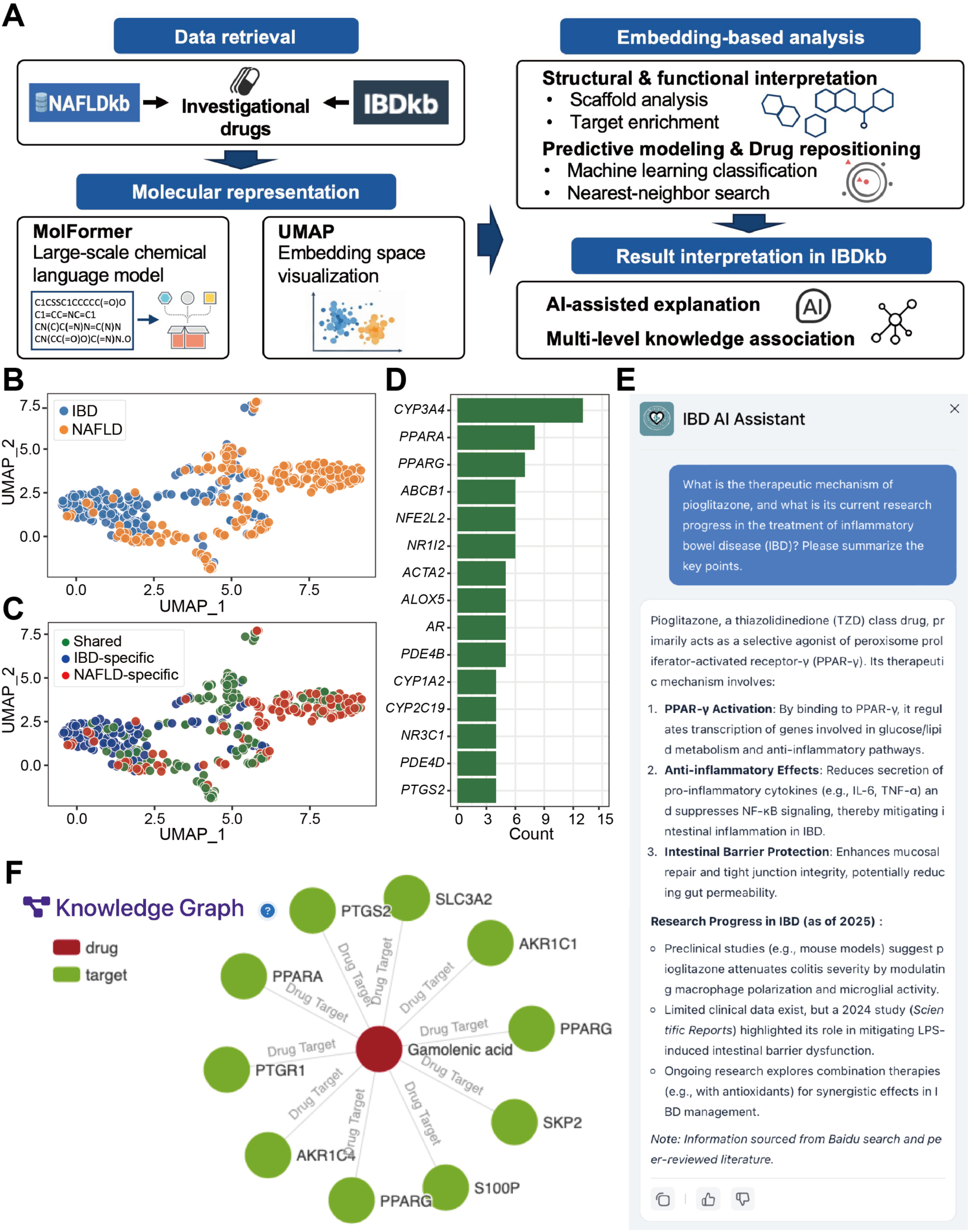
Structure-aware drug comparison and repositioning. (A) The workflow of structure-aware comparative analysis of investigational drugs for IBD and NAFLD based on molecular embeddings. (B) UMAP visualization of MolFormer-derived molecular embeddings. (C) UMAP projection of the same embedding space annotated by Murcko scaffold categories, including IBD-specific scaffolds, NAFLD-specific scaffolds, and scaffolds shared between the two disease indications. (D) Target distribution of drugs associated with the shared scaffold category. Bar plots show the top 15 most frequently occurring targets. (E) Results retrieved using the global AI assistant to query the therapeutic mechanisms of pioglitazone and its current research progress in IBD. (F) Knowledge association network displayed on the detail page of gamolenic acid.

By transforming heterogeneous drug information into unified embedding representations, we generated AI-ready datasets that support direct integration with downstream machine learning tasks, including classification, similarity analysis, and knowledge discovery. To interpret the structural basis, Murcko scaffold analysis was performed, categorizing scaffolds into IBD-specific, NAFLD-specific, and shared groups (Table S1). These categories exhibited distinct and coherent spatial distributions in the embedding space (Figure 5C). Target enrichment analysis revealed that shared scaffolds were enriched for targets involved in drug metabolism/transport, inflammation, and immune signaling, such as PPARs, suggesting potential mechanistic overlap (Figure 5D). Building on these embedding features, five machine-learning classifiers were trained to distinguish IBD drugs from NAFLD drugs. All models achieved robust performance, with 5-fold cross-validation mean area under the curve (AUC) values exceeding 0.93 (Figure S2A), further confirming structural differences.

Nearest-neighbor analysis in the embedding space explored cross-disease drug repositioning opportunities. Several NAFLD drugs (e.g., LYS006, PF-05221304, pioglitazone) were identified as closest structural neighbors to multiple IBD drugs, suggesting possible therapeutic transferability (Figure S2B, Table S2). Conversely, some IBD drugs (e.g., 2’-fucosyllactose, probiotic, gamolenic acid) showed high structural similarity to NAFLD agents (Figure S2C, Table S3). This analysis identifies structural proximity as a hypothesis-generating signal, whereas pharmacological similarity and clinical translatability require further validation.

We further leveraged IBDkb’s AI tools to interpret the results. Pioglitazone, structurally proximal to IBD drugs (Figure S2B), was characterized as a PPARγ agonist with anti-inflammatory and intestinal barrier-protective effects (Figure 5E) [29]. A global search in IBDkb identified 15 publications on pioglitazone in IBD (Figure S2D). Although PPARγ has emerged as a novel therapeutic target in IBD [30], existing studies are largely preclinical [31]. In addition, 90 bioactive compound records associated with pioglitazone were retrieved (Figure S2D, Table S4), providing potential insights for IBD drug development. Gamolenic acid, structurally proximal to NAFLD drugs (Figure S2C), was linked to 10 drug target records in IBDkb, including PPARs (Figure 5F), highlighting its potential relevance in NAFLD. This case study demonstrates how IBDkb, integrated with molecular representation models, can facilitate structure-aware drug comparison and support cross-disease repositioning hypotheses.

## Discussion

IBDkb is a comprehensive, therapy-oriented knowledge base and platform for IBD, designed to systematically organize and connect heterogeneous information across drugs, therapeutic strategies, experimental models, clinical trials, and literature. While existing IBD databases predominantly focus on omics-level data [12–19], their systematic integration with therapeutic development remains limited. In contrast, IBDkb adopts a drug development-centered organizational framework that explicitly linking mechanistic insights with therapeutic strategies and clinical evidence, thereby supporting translational research and data-driven therapeutic discovery.

Despite advances in understanding IBD pathophysiology, the gap between mechanistic insights and clinical translation contributes to high preclinical attrition rates in IBD drug development, for which artificial intelligence-based large-scale data analysis offers promising opportunities to improve efficiency [32]. By harmonizing drug-related information, mechanistic annotations, and clinical trial evidence within a unified framework, IBDkb enables systematic comparisons across therapeutic strategies and disease contexts. This integrative design facilitates applications such as drug repurposing and mechanism-driven trial design, while providing structured and computable background knowledge for AI-assisted drug discovery workflows. IBDkb further offers downloadable AI-ready datasets that enable direct application of machine learning models for drug classification, similarity analysis, and knowledge discovery, lowering the technical barrier for AI-based analyses.

The incorporation of AI-assisted knowledge exploration further enhances IBDkb’s utility. With the rapid advancement of large language models, AI-based tools have demonstrated substantial potential for biomedical knowledge synthesis and reasoning [33]. Retrieval-augmented generation (RAG) frameworks, combined with curated domain knowledge, are increasingly applied to tasks such as literature interpretation and hypothesis generation [34]. Compared with traditional query-based databases, the AI assistant embedded within IBDkb enables more flexible and efficient exploration of complex therapeutic evidence, facilitates connections between drug mechanisms and clinical outcomes, and supports hypothesis generation through context-aware interactions.

Despite these strengths, IBDkb also has limitations. The current version primarily relies on published studies and publicly clinical trial data, necessitating continuous updates. Future extensions will aim to incorporate longitudinal treatment outcomes and multi-omics data associated with therapeutic responses. From an AI perspective, further integration with predictive modeling frameworks and digital twin-based approaches may enable more advanced simulation, outcome prediction, and personalized therapeutic exploration. These developments will further strengthen IBDkb as a dynamic and extensible platform for IBD research and drug development.

## Conclusion

IBDkb is a unique, community-oriented resource that integrates broad, curated IBD data with intuitive visualization and cutting-edge AI-enhanced tools. Freely available without registration, it supports downloadable datasets and welcomes user feedback. By bridging molecular insights with therapeutic development, IBDkb aims to accelerate precision medicine and innovation in IBD research.

## Acknowledgments

We gratefully acknowledge the following sources of financial support: the National Natural Science Foundation of China (Grant Nos. 82170542 to R.Z., 82304351 to L.L., 32200529 and 32570768 to D.W., and 32470098 to N.J.), the Shanghai Oriental Talents Program (Grant BJJY2024098 to R.Z.), the China Postdoctoral Science Foundation (Grant 2025M772680 to L.L.), and the Doctoral Student Special Project of the Young Talent Support Program by the China Association for Science and Technology (CAST) (to W.C.). The funders had no role in the study design, data collection and analysis, decision to publish, or preparation of the manuscript. We also thank the Center for Scientific Computing and the Supercomputing Center of the School of Life Sciences and Technology, Tongji University, for providing computational resources and support.

## Authors’ contributions

D.W., R.Z., and P.X. conceived and designed the project. Each author has contributed significantly to the submitted work. L. T. and S.S. collected and analyzed the data. L.T. and S.S. developed and tested code for data processing and analysis. L.T., S.S., Z.L., and B.Y. were responsible for website development, testing, and maintenance. L.T. drafted the manuscript and prepared figures. All authors contributed to the interpretation of the results and critical revision of the manuscript for important intellectual content. All authors have read and approved the final manuscript.

## Availability of data and materials

All the software packages used in this study are open-source and publicly available. All code used and/or analyzed in this study are available on GitHub at https://github.com/tjcadd2020/IBDkb.

## Reference

1. Graham DB, Xavier RJ: Pathway paradigms revealed from the genetics of inflammatory bowel disease. Nature 2020, 578:527–539.

2. Le Berre C, Honap S, Peyrin-Biroulet L: Ulcerative colitis. Lancet 2023, 402:571–584.

3. Dolinger M, Torres J, Vermeire S: Crohn’s disease. Lancet 2024, 403:1177–1191.

4. Porter RJ, Arends MJ, Churchhouse AMD, Din S: Inflammatory Bowel Disease-Associated Colorectal Cancer: Translational Risks from Mechanisms to Medicines. Journal of Crohn’s and Colitis 2021, 15:2131–2141.

5. Hracs L, Windsor JW, Gorospe J, Cummings M, Coward S, Buie MJ, Quan J, Goddard Q, Caplan L, Markovinović A, et al: Global evolution of inflammatory bowel disease across epidemiologic stages. Nature 2025, 642:458–466.

6. Kaplan GG: The global burden of inflammatory bowel disease: from 2025 to 2045. Nature Reviews Gastroenterology & Hepatology 2025, 22:708–720.

7. Chang JT: Pathophysiology of Inflammatory Bowel Diseases. New England Journal of Medicine 2020, 383:2652–2664.

8. Vanhove W, Nys K, Vermeire S: Therapeutic innovations in inflammatory bowel diseases. Clin Pharmacol Ther 2016, 99:49–58.

9. Cannarozzi AL, Latiano A, Massimino L, Bossa F, Giuliani F, Riva M, Ungaro F, Guerra M, Brina ALD, Biscaglia G, et al: Inflammatory bowel disease genomics, transcriptomics, proteomics and metagenomics meet artificial intelligence. United European Gastroenterol J 2024, 12:1461–1480.

10. Zhang Y, Thomas JP, Korcsmaros T, Gul L: Integrating multi-omics to unravel host-microbiome interactions in inflammatory bowel disease. Cell Reports Medicine 2024, 5:101738.

11. Chen W, Li Y, Wang W, Gao S, Hu J, Xiang B, Wu D, Jiao N, Xu T, Zhi M, et al: Enhanced microbiota profiling in patients with quiescent Crohn’s disease through comparison with paired healthy first-degree relatives. Cell Reports Medicine 2024, 5:101624.

12. Rivas MA, Avila BE, Koskela J, Huang H, Stevens C, Pirinen M, Haritunians T, Neale BM, Kurki M, Ganna A, et al: Insights into the genetic epidemiology of Crohn’s and rare diseases in the Ashkenazi Jewish population. PLoS Genet 2018, 14:e1007329.

13. Lloyd-Price J, Arze C, Ananthakrishnan AN, Schirmer M, Avila-Pacheco J, Poon TW, Andrews E, Ajami NJ, Bonham KS, Brislawn CJ, et al: Multi-omics of the gut microbial ecosystem in inflammatory bowel diseases. Nature 2019, 569:655–662.

14. Kai N, Qingsong C, Kejia M, Weiwei L, Xing W, Xuejie C, Lixia C, Minzi D, Yuanyuan Y, Xiaoyan W: An Inflammatory Bowel Diseases Integrated Resources Portal (IBDIRP). Database 2024, 2024.

15. Shen J, Mao AP, Zhu MM, Zhao P, Xu JJ, Zuo Z: Ulcerative Colitis Database: An Integrated Database and Toolkit for Gene Function and Medication Involved in Ulcerative Colitis. Inflamm Bowel Dis 2015, 21:1872–1882.

16. Khan F, Radovanovic A, Gojobori T, Kaur M: IBDDB: a manually curated and text-mining-enhanced database of genes involved in inflammatory bowel disease. Database (Oxford) 2021, 2021.

17. Velissari R, Ilieva M, Dao J, Miller HE, Madsen JH, Gorodkin J, Aikawa M, Ishii H, Uchida S: Systematic analysis and characterization of long non-coding RNA genes in inflammatory bowel disease. Brief Funct Genomics 2024, 23:395–405.

18. Avram V, Yadav S, Sahasrabudhe P, Chang D, Wang J: IBDTransDB: a manually curated transcriptomic database for inflammatory bowel disease. Database (Oxford) 2024, 2024.

19. Nie H, Lin P, Zhang Y, Wan Y, Li J, Yin C, Zhang L: Single-cell meta-analysis of inflammatory bowel disease with scIBD. Nature Computational Science 2023, 3:522–531.

20. Ripa JD, Ali S, Field M, Smithson J, Wangchuk P: From AI-Assisted In Silico Computational Design to Preclinical In Vivo Models: A Multi-Platform Approach to Small Molecule Anti-IBD Drug Discovery. Pharmaceuticals (Basel) 2025, 18.

21. Knox C, Wilson M, Klinger CM, Franklin M, Oler E, Wilson A, Pon A, Cox J, Chin NEL, Strawbridge SA, et al: DrugBank 6.0: the DrugBank Knowledgebase for 2024. Nucleic Acids Res 2024, 52:D1265–d1275.

22. Kim S, Chen J, Cheng T, Gindulyte A, He J, He S, Li Q, Shoemaker BA, Thiessen PA, Yu B, et al: PubChem 2023 update. Nucleic Acids Res 2023, 51:D1373–d1380.

23. Zdrazil B, Felix E, Hunter F, Manners EJ, Blackshaw J, Corbett S, de Veij M, Ioannidis H, Lopez DM, Mosquera JF, et al: The ChEMBL Database in 2023: a drug discovery platform spanning multiple bioactivity data types and time periods. Nucleic Acids Res 2024, 52:D1180–d1192.

24. Amberger JS, Bocchini CA, Scott AF, Hamosh A. OMIM.org: leveraging knowledge across phenotype-gene relationships. Nucleic Acids Res 2019, 47:D1038–d1043.

25. Zhou Y, Zhang Y, Zhao D, Yu X, Shen X, Zhou Y, Wang S, Qiu Y, Chen Y, Zhu F: TTD: Therapeutic Target Database describing target druggability information. Nucleic Acids Res 2024, 52:D1465–d1477.

26. UniProt: the Universal Protein Knowledgebase in 2025. Nucleic Acids Res 2025, 53:D609–d617.

27. Xu T, Gao W, Zhu L, Chen W, Niu C, Yin W, Ma L, Zhu X, Ling Y, Gao S, et al: NAFLDkb: A Knowledge Base and Platform for Drug Development against Nonalcoholic Fatty Liver Disease. Journal of Chemical Information and Modeling 2024, 64:2817–2828.

28. Ross J, Belgodere B, Chenthamarakshan V, Padhi I, Mroueh Y, Das P: Large-scale chemical language representations capture molecular structure and properties. Nature Machine Intelligence 2022, 4:1256–1264.

29. Zhou Z, Jin R, Gu Y, Ji Y, Lou Y, Wu J: Therapeutic Targeting of PPARγ in Nonalcoholic Fatty Liver Disease: Efficacy, Safety, and Drug Development. Drug Des Devel Ther 2025, 19:7293–7319.

30. Dubuquoy L, Rousseaux C, Thuru X, Peyrin-Biroulet L, Romano O, Chavatte P, Chamaillard M, Desreumaux P: PPARgamma as a new therapeutic target in inflammatory bowel diseases. Gut 2006, 55:1341–1349.

31. da Rocha GHO, Loiola RA, de Paula-Silva M, Shimizu F, Kanda T, Vieira A, Gosselet F, Farsky SHP: Pioglitazone Attenuates the Effects of Peripheral Inflammation in a Human In Vitro Blood-Brain Barrier Model. Int J Mol Sci 2022, 23.

32. Honap S, Jairath V, Danese S, Peyrin-Biroulet L: Navigating the complexities of drug development for inflammatory bowel disease. Nat Rev Drug Discov 2024, 23:546–562.

33. Lee P, Bubeck S, Petro J. Benefits, Limits, and Risks of GPT-4 as an AI Chatbot for Medicine. N Engl J Med 2023, 388:1233–1239.

34. Jeong M, Sohn J, Sung M, Kang J: Improving medical reasoning through retrieval and self-reflection with retrieval-augmented large language models. Bioinformatics 2024, 40:i119–i129.

