## Supplementary material for "IBDkb: an AI-enhanced integrative knowledge base for inflammatory bowel disease research and drug discovery": Figure S

### Supplementary Figures

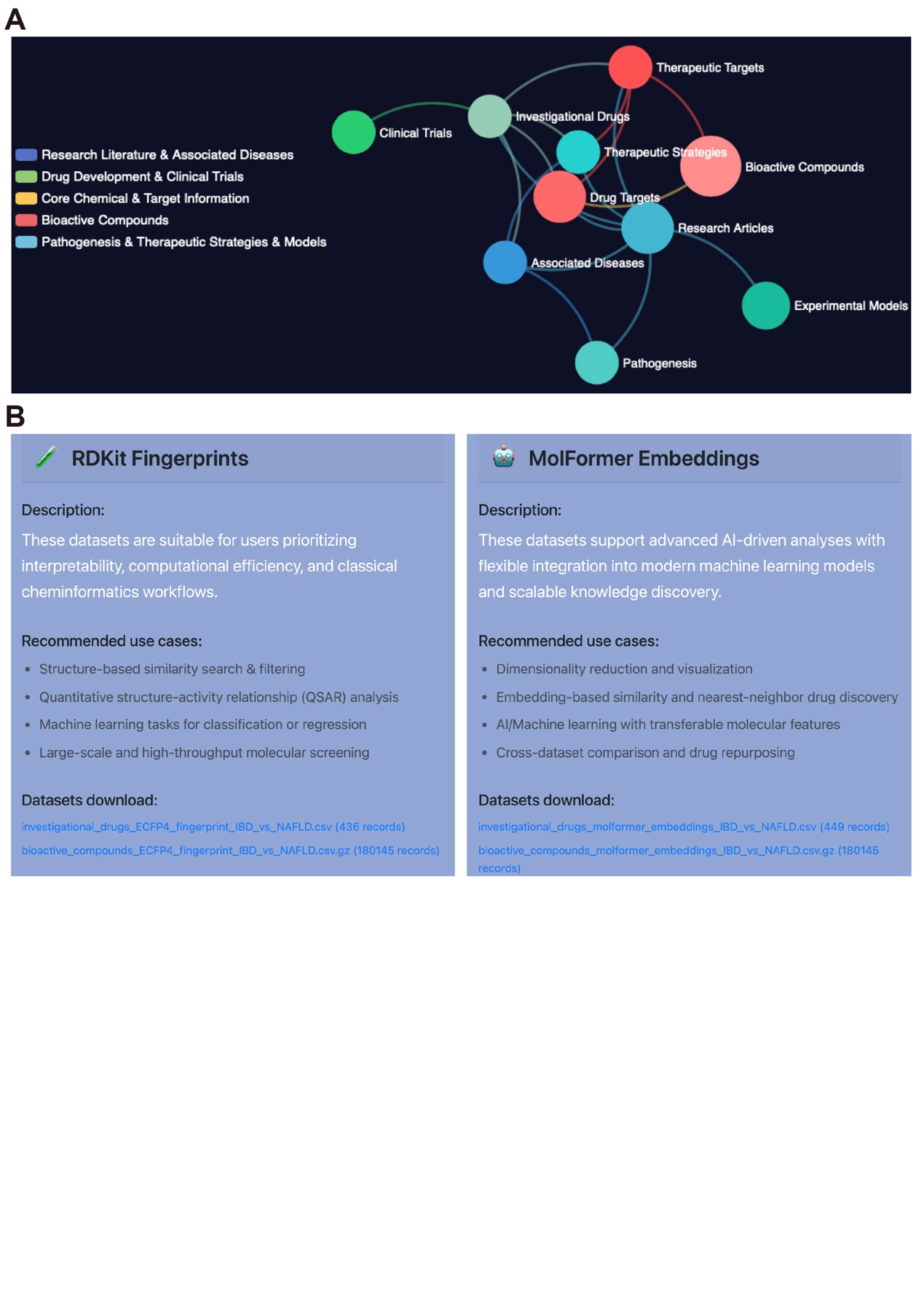

Figure S1. **Utility of the IBDkb platform.** (A) Association network illustrating the relationships among different modules within IBDkb. (B) Data resources available in the Downloads module of IBDkb.

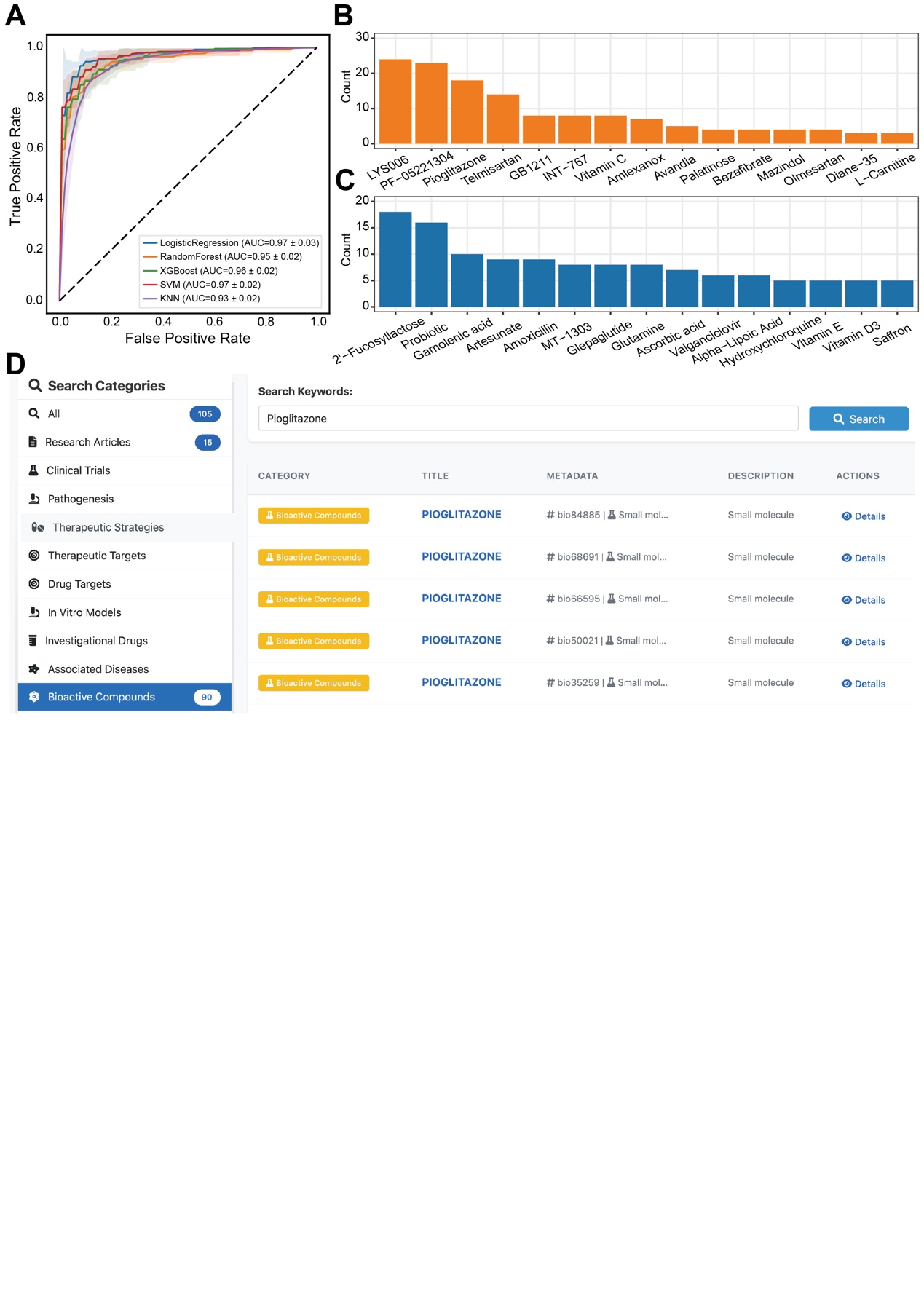

Figure S2. **Structure-aware drug comparison and repositioning**. (A) Receiver operating characteristic (ROC) curves of five machine-learning classifiers trained using molecular embedding features to discriminate IBD drugs from NAFLD drugs. Mean AUC with standard deviation of stratified five-fold cross-validation were shown. (B) Frequency of NAFLD drugs identified as the nearest neighbors for individual IBD drugs in the embedding space. Bar plots display the top 15 NAFLD drugs with the highest occurrence counts. (C) Frequency of IBD drugs identified as the nearest neighbors for individual NAFLD drugs in the embedding space. Bar plots display the top 15 IBD drugs with the highest occurrence counts. (D) Overview of the global search results for pioglitazone in IBDkb.

### Supplementary Table Legends

Table S1. MolFormer-based embedding results of investigational drugs.

Table S2. Results of the nearest-neighbor analysis from IBD to NAFLD.

Table S3. Results of the nearest-neighbor analysis from NAFLD to IBD.

Table S4. Bioactive compound records associated with pioglitazone retrieved from the global search in IBDkb.
